# Complex harmonics reveal low-dimensional manifolds of critical brain dynamics

**DOI:** 10.1101/2024.06.15.599165

**Authors:** Gustavo Deco, Yonatan Sanz Perl, Morten L. Kringelbach

## Abstract

The brain needs to perform time-critical computations to ensure survival. A potential solution lies in the non-local, distributed computation at the whole-brain level made possible by criticality and amplified by the rare long-range connections found in the brain’s unique anatomical structure. This non-locality can be captured by the mathematical structure of Schrödinger’s wave equation, which is at the heart of the novel CHARM (Complex Harmonics decomposition) framework that performs the necessary dimensional manifold reduction able to extract non-locality in critical spacetime brain dynamics. Using a large neuroimaging dataset of over 1,000 people, CHARM captured the critical, non-local/long-range nature of brain dynamics and the underlying mechanisms were established using a precise whole-brain model. Equally, CHARM revealed the significantly different critical dynamics of wakefulness and sleep. Overall, CHARM is a promising theoretical framework for capturing the low-dimensionality of the complex network dynamics observed in neuroscience and provides evidence that networks of brain regions rather than individual brain regions are the key computational engines of critical brain dynamics.

## Introduction

There is a deep conundrum at the heart of human cognition, namely how the surprisingly slow information transfer between neurons (with typical latencies of around 10-20 milliseconds) is able to solve the time-critical computational problems ensuring survival (Itoh *et al*., 2022). Paradoxically, the wetware of the brain is still better at solving problems than much faster silicon-based computers. This raises the unsolved physical problem of how the brain overcomes the limitations of speed for information transfer across spacetime.

Beyond local computation, the solution has been proposed to be time-critical distributed computation (Deco *et al*., 2023). The high dimensional space obtained from whole brain neuroimaging using functional MRI or EEG/MEG has been described by lower dimensional, spatially distributed ‘resting state networks’ following an avalanche of important research over the last decades (Biswal *et al*., 1995; Biswal *et al*., 2010; Buckner *et al*., 2008; Finn, 2021; Smith *et al*., 2009). Importantly, the computation at the network level emerges from the existence of critical dynamics producing long-range interactions which give rise to low-dimensional manifolds (Meisel *et al*., 2012; Ponce-Alvarez *et al*., 2023; Tagliazucchi *et al*., 2016; Yu *et al*., 2013). This is amplified by the long-range interactions in critical dynamics through the existence of the unique mammalian architecture with rare long-range connections between distant brain regions, which are exceptions to the predominant short-range wiring with an exponential drop off in strength over distance (Ercsey-Ravasz *et al*., 2013; Gămănuţ *et al*., 2018; Theodoni *et al*., 2020). This Bauplan with weight-distance relations supplemented with rare long-range connections between brain regions likely makes brain architecture unique among known physical systems (Deco *et al*., 2021). Thus, the underlying low-dimensional networks expressing these non-local effects and performing the necessary time-critical computations are key to understanding human cognition (Deco *et al*., 2023).

Here, we developed CHARM (Complex Harmonics decomposition) to provide the low-dimensional manifold reduction able to capture non-locality in critical spacetime brain dynamics. The CHARM framework uses the formal mathematical definition of the manifold reduction problem (Atasoy *et al*., 2016; Belkin and Niyogi, 2003; Gilpin, 2024). Crucially, however, instead of deriving this from the heat equation which is solely conserving local neighbourhood, we derived this from the Schrödinger equation to produce a complex kernel able to capture the long-range, non-local effects of critical brain dynamics (Schleich *et al*., 2013; Schrödinger, 1926a, b).

We derive the mathematical structure of CHARM and we demonstrate the efficiency and robustness of this fundamental framework for understanding brain dynamics using large-scale neuroimaging data from over 1,000 human participants. We show that this mathematical structure captures the critical, non-local/long-range nature of brain dynamics, and that this is significantly better than the best competing method. We establish the mechanistic reasons for why this mathematical formalism is able to capture the long-range, non-local critical interactions in brain dynamics by using a precise whole-brain model. Finally, we show how CHARM can reveal the significant differences in critical dynamics of different brain states of wakefulness and sleep.

### Complex Harmonic decomposition (CHARM) of brain dynamics

We present here the mathematical formulation of the novel Complex Harmonic decomposition (CHARM) framework for manifold reduction of complex dynamical systems such as the brain. **Figure 1A** shows an overview of how CHARM can be used to decompose the complex high dimensional space neuroimaging data of whole-brain dynamics into low-dimensional manifold networks. In **Figure 1B** we show a cartoon of the process of decomposition using classic harmonics and CHARM on a hidden underlying manifold with colourmaps indicating neighbourhood and the lines indicating long-range connections. We then use whole-brain modelling to elucidate the underlying mechanisms of CHARM to capture the two aspects of non-locality in brain dynamics, namely criticality and the role of anatomical long-range connections (**Figure 1C**).

**Figure 1.**
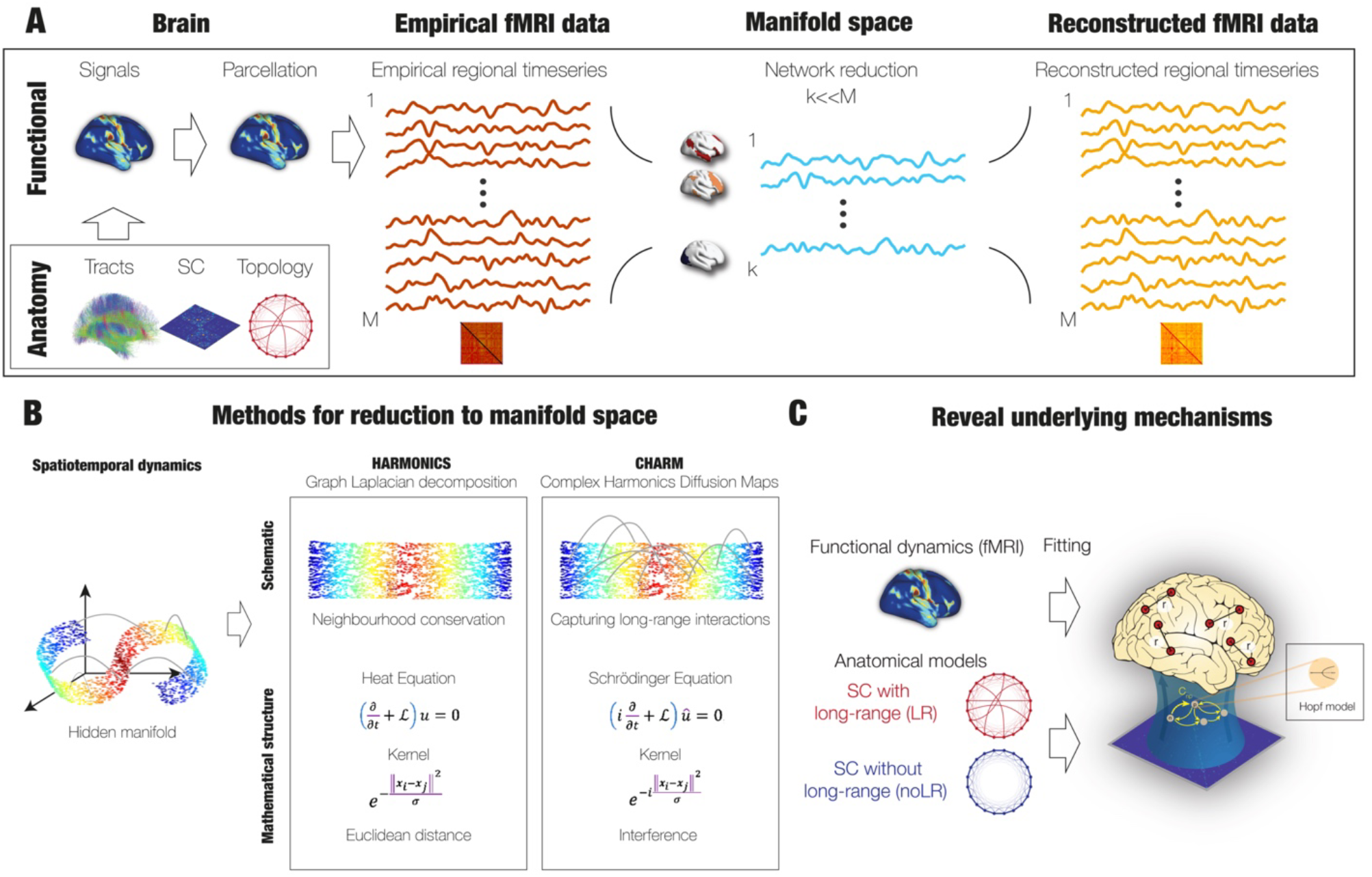
Extraction of low-dimensional manifold of networks in brain dynamics. **A)** Neuroimaging data of whole-brain dynamics yields a higher dimensional space, which can be decomposed into low-dimensional manifold networks. The brain data is measured with functional MRI and arises from the neural activity coupled through the underlying anatomy with a special topology. Uniquely to mammalian brain architecture there are long-range (LR) connections, which are exceptions to short-range wiring with an exponential drop off in strength over distance, called the exponential distance rule (EDR). The functional activity is divided into specific brain regions in a given parcellation with M regions. In red, we show the M high-dimensional source space with examples of the individual timecourses and the M × M functional connectivity source matrix. This can be decomposed into low-dimensional manifolds, given the existence of critical dynamics producing long-range interactions which in the brain are further amplified by the anatomical LR connectivity. The k low-dimensional manifold space is shown in light blue with the timeseries activity in the underlying manifold networks. The quality of this manifold space can be measured by comparing the M high-dimensional reconstructed space, here shown in orange with examples of the individual timecourses and the M × M functional connectivity reconstructed matrix. **B)** The panel is a cartoon of the process of decomposition using two different methodologies on a hidden underlying manifold with colourmaps indicating neighbourhood and the lines indicating long-range connections (far left). The first box shows illustrates the principles underlying harmonics graph Laplacian decomposition. Thanks to the Gaussian kernel derived from the Heat Equation, this conserves neighbourhood relationships as reflected in the ability of the method to decompose the manifold into a low dimensional space (where the colouring matches the neighbourhood of the hidden manifold). The second box illustrates the CHARM (Complex Harmonics Diffusion Maps) framework. This uses a complex kernel derived from the Schrödinger Wave equation able to capture not only the neighbourhood relationships but also the non-local long-range interactions, thanks to the interference in the non-local brain dynamics brought about by the underlying criticality amplified by the anatomical long-range connectivity. **C)** Whole-brain modelling was used to test the relevance of harmonics and CHARM to capture the two aspects of non-locality in brain dynamics, namely criticality and the role of anatomical long-range connections.

CHARM is used on complex neuroimaging data where whole-brain activity is measured using fMRI which produces BOLD timeseries for over 1,000 healthy human participants. In order to reduce the dimensionality of these timeseries, we define ***x***_*i*_ ∈ ℛ^*M*^, which denotes the column vector containing the BOLD signal of the *M* brain regions at the *i*-th time point of the time series. The matrix of all brain regions can then be defined by ***X*** = [***x***_1_, ***x***_2_, …, ***x***_*N*_] ∈ ℛ^*M*×*N*^, where the columns of the matrix span a time window of *N* time point observations. We assume that the brain dynamics lie on a sufficiently smooth low-dimensional (say of dimension *k* ≪ *M*) manifold which is embedded in the high-dimensional ℛ^*M*^ space. With this notation, the generic problem of dimensionality reduction can be defined as follows: Given a set ***X***, find a set of points ***Y*** = [***y***_1_, ***y***_2_, …, ***y***_*N*_] ∈ *ℛ*^*k*×*N*^such that ***y***_*i*_ ∈ ℛ^*k*^ “represents” ***x***_*i*_.

The formulation of manifold reduction in continuum space allows for the analytical derivation in discrete space (Belkin and Niyogi, 2003). In continuum space, the manifold reduction can be formally defined as follows: Let ℳ be a smooth, compact, *m*-dimensional Riemannian source space (Rosenberg, 1997). First consider the one-dimensional reduction, where the manifold is a real line which is defined in such way that the points close together in the source space are also mapped close together on the manifold line. Let *f* be such a map from source space to a manifold line, *f*: ℳ → ℛ^1^ which is twice differentiable. Belkin and Niyogi have shown that this map can be found by minimizing the cost function 𝔥

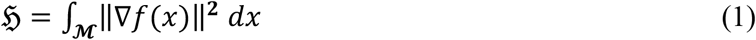

Minimising the objective function 𝔥 is equivalent to finding the eigenfunctions of the Laplace Beltrami operator ℒ, defined by ℒ*f* = −*div*∇(*f*). Let us denote by *f*_*i*_ the eigenfunctions of ℒ. Consequently, the map to a line is defined by the first non-trivial eigenfunction. Please note that the first eigenfunction is a trivial constant function that maps the entire manifold to a single point. In the general case the map defining the dimensionality reduction conserving neighbourhood is given by

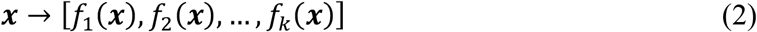

The Laplace Beltrami operator on differentiable functions on a source space ℳ is intimately related to heat flow as shown by Belkin and Niyogi (Belkin and Niyogi, 2003), who derived the harmonic decomposition from the heat equation, i.e. the partial differential equation 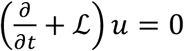. They showed that the solution is given by *u*(***x***, *t*) = ∫_ℳ_ *H*_*t*_(***x, y***)*f*(***y***) *d****y*** where *H*_*t*_(***x, y***) is the heat kernel, the Green’s function for this partial differential equation. In other words, the initial heat distribution *u*(***x***, 0) = *f*(***x***). Therefore,

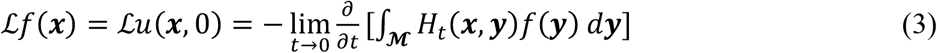

Given that the kernel of the heat equation is approximately Gaussian, this leads to

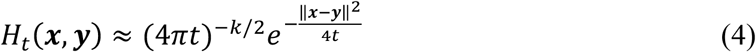

Notice that as *t* tends to 0, the heat kernel *H*_*t*_(*x, y*) tends to Dirac’s *δ*-function, that is, 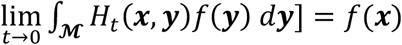. Therefore, for small *t* from the definition of the derivative, we have

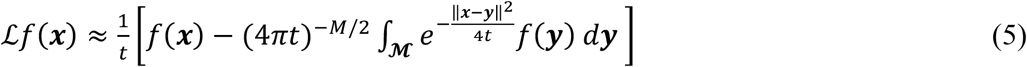

If [***x***_1_, ***x***_2_, …, ***x***_*N*_] are data points on ℳ, the last expression can be approximated by

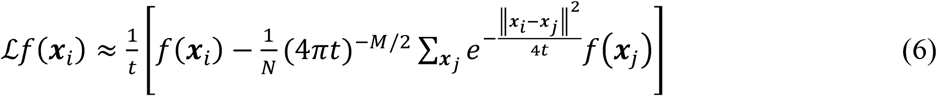

Given that global coefficients will not affect the eigenvectors of the discrete Laplacian, we can conveniently rescale the last equation such that the eigenfunctions of ℒ can be discretised by

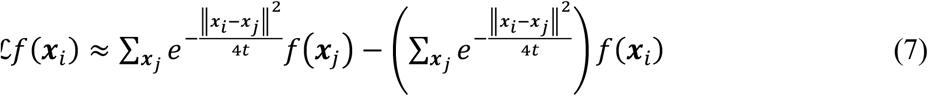

Which determines the discretisation precisely as the graph Laplacian matrix as shown by Belkin and Niyogi. The coordinates of the nonlinear projection of a point ***x***_*i*_ in the reduced manifold latent *k* dimensional space spanned by the first *k* eigenvectors [***φ***_1_, ***φ***_2_, …, ***φ***_*k*_] of the graph Laplacian is given by:

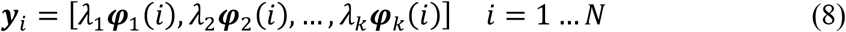

Here ***φ***_*j*_(*i*) denotes the *i*-th element of the eigenvector ***φ***_*j*_. The embedding dimension can be determined by the spectral gap in the eigenvalues of the final decomposition, which provides the classic harmonic decomposition. To sum up, the origin of the Gaussian kernel can be analytically derived from the definition of the manifold reduction problem in the continuum space by solving the heat equation when the time tends to zero. A direct interpretation of the Gaussian kernel functioning is thus as the implementation of manifold reduction conserving the discrete neighbourhood space, which is the basis of harmonic decomposition in **Equation 8**.

However, given that brain dynamics are not just local, we were inspired by Schrödinger to develop a complex kernel to capture the long-range functional connectivity of the human brain, which has been shown to improve information transfer (Deco *et al*., 2023; Deco *et al*., 2021). This complex kernel should be able to capture the interference and aggregation of information transfer mediated by long-range interactions between brain regions. This would capture the effect of long-range dependencies essential in brain dynamics due both to the role by long-range anatomical exceptions but also to the dynamical criticality causing the emergence of long-range functional correlations.

Here, we first provide a formal mathematical derivation. The key idea is to find an alternative route to capture the non-local effects by finding a complex equation with time going to zero from the imaginary axis, rather than just have time go to zero from the real axis (as was the case for the heat equation). Specifically, beyond the heat equation, which is only defined in real space, we propose to use the Schrödinger equation (Schrödinger, 1926a), i.e., the partial differential equation 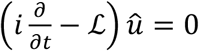, which corresponds to the case of a free particle. Here we simplified Schrödinger’s equation by using the constants 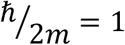. The solution is given by 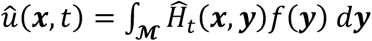, where 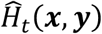 is the free particle Schrödinger kernel, also known as Green’s function for this partial differential equation. This allows us to carry out exactly the same steps as above but now starting from Schrödinger’s equation instead of the heat equation. The results are equivalent and easy to reproduce using the Wick transformation *t* → *it*, arriving to the equivalent convenient discretisation of the eigenfunctions of ℒ, from the Schrödinger perspective, as

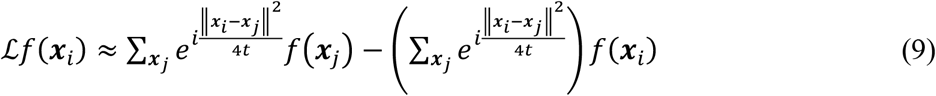

In other words, we now get an approximation of the graph Laplacian matrix but with a new complex kernel for the matrix 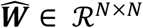, whose elements 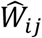 are given by

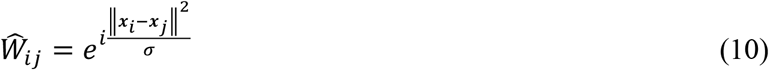

where ‖ ‖ computes the distance between two points using the Euclidean L2 norm, and σ serves as a scale parameter of the kernel. Instead of the graph Laplacian, it is more convenient to work with the transition probability matrix. In order to best capture long-range/non-local interactions, which produces constructive and destructive interferences, we used a two-step procedure: First, we define the *t*-steps diffusion matrix by taking the power *t* of the diffusion matrix 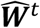 . Second, similar to Schrödinger (Schleich *et al*., 2013; Schrödinger, 1982), we define the non-normalised probability transition matrix by the square module of the *t*-steps diffusion matrix:

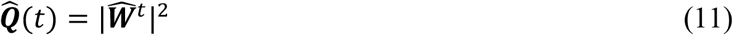

where | | notates the module. We define the corresponding diagonal normalisation matrix 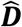 with elements as follows

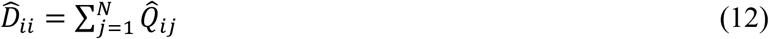

where the normalised transition probability matrix 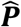 is given by

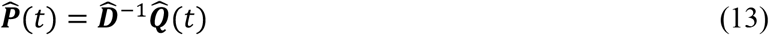

Applying singular value decomposition on 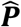, we get

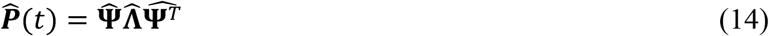

with 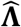 being the diagonal matrix that stores the *M* eigenvalues (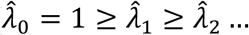, with the first being the trivial eigenvalue that is equal to 1 given that ***P*** is a Markovian matrix) and 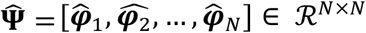 whose columns 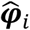 are the eigenvectors of 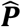.

Consequently, the CHARM manifold reduction, 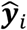, is given by the coordinates of the nonlinear projection of a point ***x***_*i*_ in the reduced manifold latent *k* dimensional space spanned by the first *k* eigenvectors 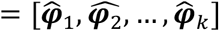 of the matrix 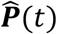

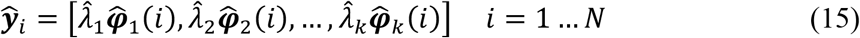

## Results

Here we tested CHARM on large-scale neuroimaging data from over 1,000 healthy human participants. First, we compared CHARM with the classic harmonic decomposition method. Second, we used whole-brain modelling to reveal the underlying mechanisms of CHARM. Finally, we showed how CHARM can reveal the significant differences in critical dynamics between the different brain states of wakefulness and sleep.

### CHARM is significantly better than harmonics for manifold reduction

We assessed the quality of the low-dimensional manifold extracted with harmonics and CHARM by comparing the reconstructed and source FC matrices. **Figure 2A** (left panels) compare harmonics and CHARM as a function of diffusion steps and a scale parameter of the kernel with the width σ. In the first row of matrices the colour scale indicates the level of correlation, while in the second row the colour scale indicates the quadratic errors between reconstructed and source FC matrices. As can be seen the optimal value of σ is indicated by a star and a box around the evolution over the diffusion steps. We zoom in on this evolution in **Figure 2A** (middle panel) which includes the dispersion over participants of the level of correlation and error as a function the diffusion step. **Figure 2A** (right panel) shows violinplots of the best level of correlation (across participants).

**Figure 2.**
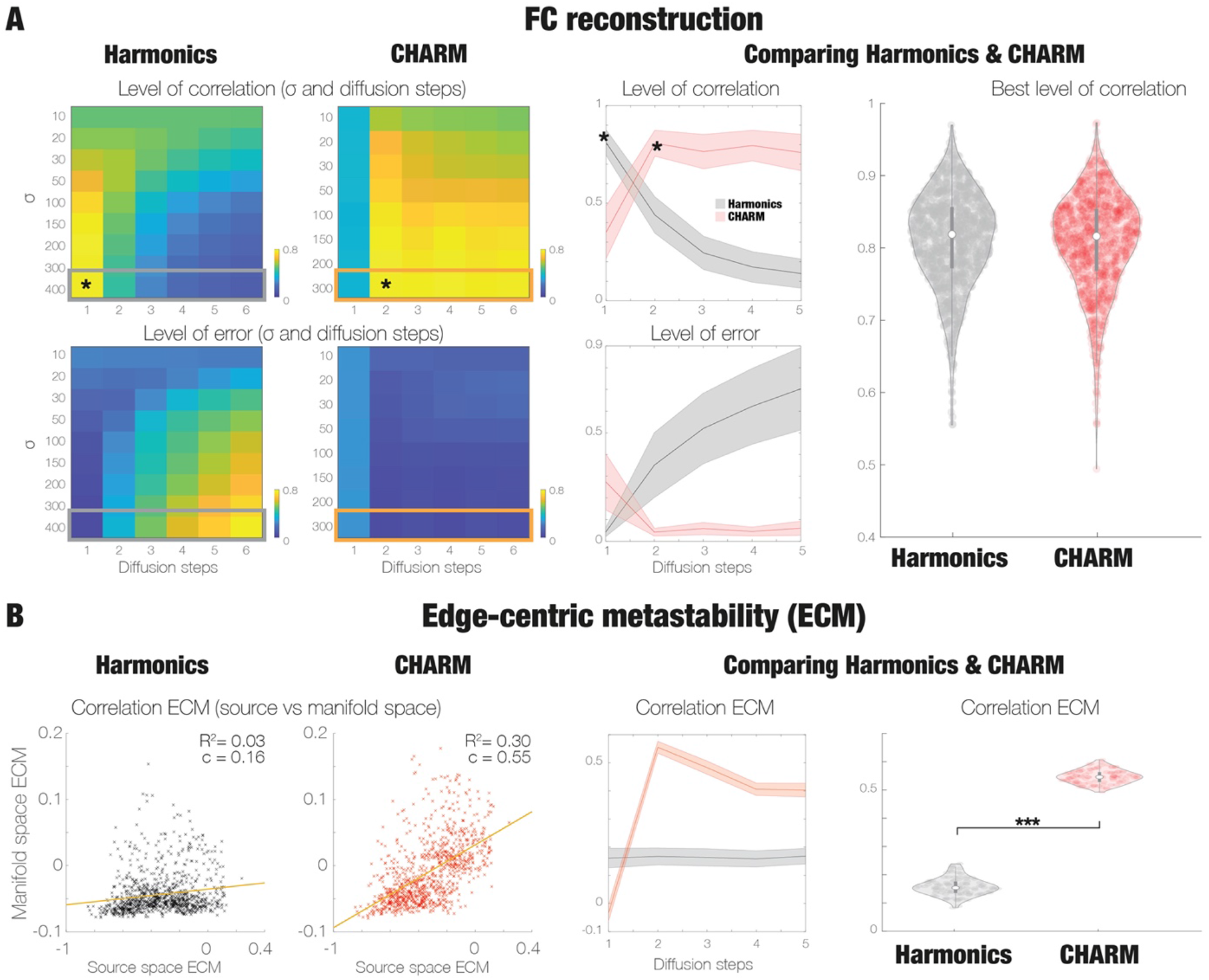
CHARM is better than harmonics for manifold reduction. **A)** The quality of the low-dimensional manifold extracted with harmonics and CHARM is assessed by comparing the reconstructed and source FC matrices. Left panels compare harmonics and CHARM as a function of diffusion steps and a scale parameter of the kernel (width σ). The colour scale in the first row of matrices indicates the correlation and the second row indicates the quadratic errors between reconstructed and source FC matrices. For each method, the optimal value of σ is indicated by a star and a box around the evolution over the diffusion steps. This evolution is shown even more clearly in the middle panel which also includes the dispersion over participants of the level of correlation and error as a function the diffusion step. The final panel shows the violinplots of the best level of correlation (across participants) for the two methods. As can be clearly seen, they both show very similar abilities in reconstructing the source signals. **B)** However, beyond the static grand average nature of FC reconstruction, we were able to measure the ability of harmonics and CHARM capture the full spacetime dynamic, for which we used the sensitive measure of edge-centric metastability (ECM). We show the scatterplots for each of them, measuring the degree of correlation across participants of ECM for the source and manifold spaces. As can be seen clearly, CHARM outperforms harmonics. Similarly, the middle panel shows the ability of each method to capture ECM as a function diffusion steps. The right panel shows the violin plots for ECM at the optimal diffusion step for each methodology. Again, as can be clearly seen from these two panels, CHARM significantly outperforms the harmonics framework, suggesting that the non-local effects play a major role in brain dynamics (p<0.001).

The results show that both methodologies show very similar abilities in reconstructing the source signals. Yet, beyond the static grand average nature of FC reconstruction, we used the highly sensitive measure of edge-centric metastability (ECM) to capture the full spacetime dynamic (see *Methods*). **Figure 2B** (left panels) shows scatterplots for harmonics and CHARM measuring the degree of correlation across participants of ECM for the source and manifold spaces. CHARM significantly outperforms the other two methodologies. **Figure 2B** (middle panel) shows the ability of each method to capture ECM as a function diffusion steps. **Figure 2B** (right panel) shows violin plots for ECM at the optimal diffusion step for each methodology. As can be seen from the plots, CHARM significantly outperforms the harmonics framework, suggesting that the non-local effects play a major role in brain dynamics.

### Whole-brain modelling reveals mechanistic principles of CHARM

**Figure 3A** shows the principles of fitting a Hopf whole-brain model to the large-scale HCP neuroimaging data from 1,003 human participants (see Methods). In **Figure 3B**, the global coupling parameter of the whole-brain was varied to obtain optimal fitting to the empirical Kuramoto metastability (blue curve) of the whole-brain model’s Kuramoto metastability (red curve) at g=0.2. Interestingly, at this optimal working point, the error of fit to the FC (green curve) is also reaching a plateau. Importantly, the model’s Kuramoto metastability is also maximal as an excellent proxy for criticality.

**Figure 3.**
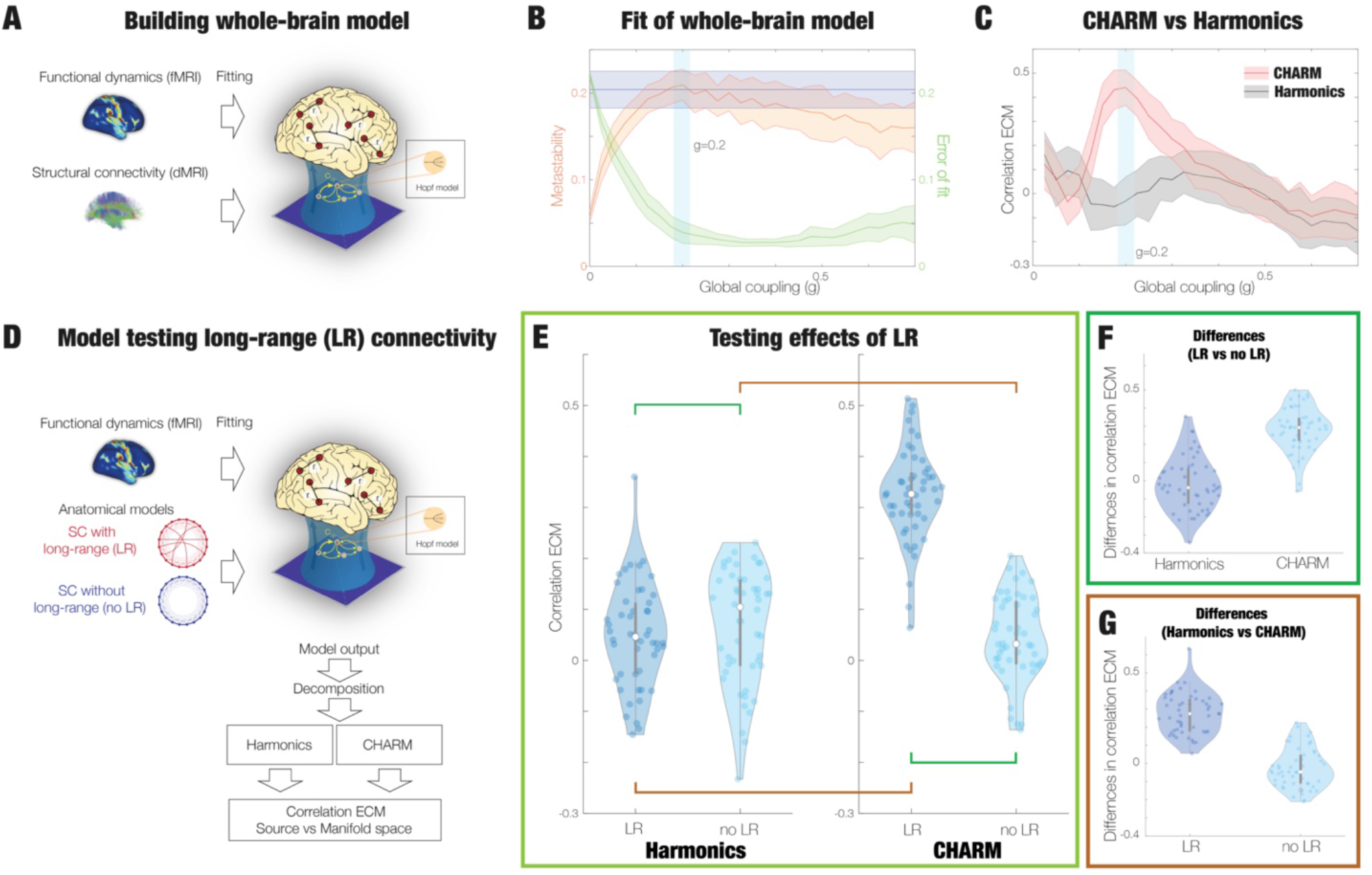
Mechanistic principles of CHARM. Using whole-brain modelling we show that CHARM outperforms harmonics in capturing non-locality in brain dynamics. **A)** We fitted a Hopf whole-brain model to the large-scale HCP neuroimaging data from 1003 participants. **B)** Varying the global coupling parameter showed optimal fitting to the empirical Kuramoto metastability (blue curve) of the whole-brain model’s Kuramoto metastability (red curve) at g=0.2. At this optimal working point, the error of fit to the FC (green curve) is also reaching a plateau. Note that at this working point, the model’s Kuramoto metastability is also maximal, which is an excellent proxy for criticality (see text). **C)** Most importantly, at the optimal working point of the whole-brain model, CHARM not only shows the largest ECM correlation between source and manifold spaces across participants but also significantly outperforms the harmonics framework. This demonstrates that CHARM is particularly well suited for extracting the non-local effects driven by the criticality. Even more, given that the best fitting of the whole-brain model is at the proxy for criticality, this reinforces the superiority of CHARM over harmonics to extract non-local critical effects in empirical brain data. **D)** In order to show the effects of the rare anatomical long-range (LR) exceptions to the general exponential distance rule of brain wiring, we constructed two models with LR (LR) and without (no LR). **E)** As can be seen, CHARM significantly outperforms harmonics in capturing the amplification of the brain dynamics by LR in terms of the ECM correlation. **F)** Reinforcing this, the figure plots the difference between LR and no LR, showing the significant difference when using CHARM. **G)** Equally, plotting the difference between harmonics and CHARM as a function LR and no LR, shows again a significant difference favouring CHARM.

But most importantly, as shown in **Figure 3C**, at the optimal working point of the whole-brain model, CHARM not only shows the largest ECM correlation between source and manifold spaces across participants but also significantly outperforms the harmonics framework. This clearly shows that CHARM is excellent in extracting the non-local effects driven by criticality. Given that the best fitting of the whole-brain model is at this proxy for criticality, this demonstrates the superiority of CHARM over harmonics for extracting non-local critical effects in empirical brain data.

**Figure 3D** shows the general framework for investigating the effects of the rare anatomical long-range (LR) exceptions to the general exponential distance rule of brain wiring. As can be seen, we constructed two models with LR (LR) and without (no LR).

The results presented in **Figure 3E** show that CHARM significantly outperforms harmonics in capturing the amplification of the brain dynamics by LR in terms of the ECM correlation. Furthermore, **Figure 3F** reinforces this by plotting the difference between LR and no LR. The results show significant difference when using CHARM compared to harmonics. Finally, as shown in **Figure 3G**, which plots the difference between harmonics and CHARM as a function LR and no LR, CHARM significantly outperforms harmonics.

### CHARM captures the distinct network interactions between different brain states

Using CHARM on data from human participants in wakefulness and deep sleep shows the excellent capability of CHARM to capture the distinct network interactions in the latent manifold space. **Figure 4A** shows the two matrices of the shifted functional connectivity between seven manifold networks reflect different interactions in wakefulness (top) and deep sleep (bottom). As can been in the small black and white matrix, there are significant differences (marked with white squares) between the two matrices.

**Figure 4.**
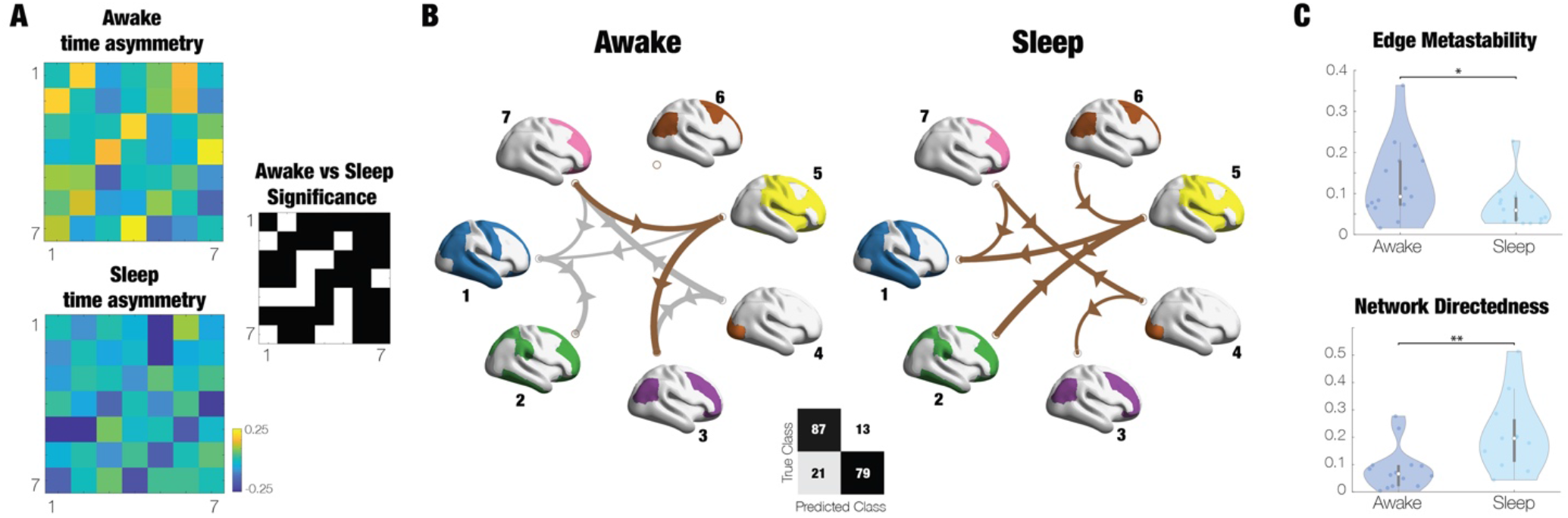
CHARM captures the distinct network interactions between different brain states. **A)** The two matrices of the shifted functional connectivity between seven manifold networks reflect different interactions in wakefulness (top) and deep sleep (bottom). Importantly, there are significant differences (white squares) between the two matrices (p<0.05). **B)** These time asymmetries are illustrated (for the top 20 percentage) of arrows between the seven manifold networks for awake vs sleep, where the grey arrows correspond to negative and brown to positive interactions, and their thickness to the value of these interactions. As can be seen the flow of interaction is very different for wakefulness compared to deep sleep. Machine learning was able to separate these interactions with an accuracy of 84 percent. **C)** The spatiotemporal characteristics in the CHARM latent space can be captured by the Edge metastability which is significantly different between awake and sleep (upper panel, p<0.05). Similarly, the reconfiguration of the hierarchical information flow is captured by the measure of network directedness of the shifted functional connectivity of the manifold network, which is also significantly different between awake and sleep (lower panel, p<0.01).

**Figure 4B** shows these time asymmetries for the top 20 percentage, illustrated by arrows between the seven manifold networks for awake vs sleep, where the grey arrows correspond to negative and brown to positive interactions, and their thickness to the value of these interactions. The flow of interaction is clearly very different for wakefulness compared to deep sleep. In fact, machine learning significantly separate these interactions with an accuracy of 84 percent (see *Methods*).

**Figure 4C** (upper panel) shows the spatiotemporal characteristics in the CHARM latent space captured by ECM, which is significantly different between awake and sleep. Similarly, the reconfiguration of the hierarchical information flow is shown in **Figure 4D** (lower panel) which captures by the measure of network directedness of the shifted functional connectivity of the manifold network. This is also significantly different between awake and sleep.

## Discussion

Recent research has shown that higher-order interactions naturally emerge from dimension reduction, providing insights into the origin of higher-order interactions in complex systems (Thibeault *et al*., 2024). Here we were inspired by the mathematical structure of Schrödinger’s wave equation to create the novel CHARM (Complex Harmonics decomposition) framework that can perform the dimensional manifold reduction able to extract non-locality in critical spacetime brain dynamics. We compared CHARM to the powerful harmonic decomposition method when used on a large neuroimaging dataset of over 1,000 people. Importantly, CHARM outperformed harmonics in terms capturing the underlying higher-order interactions in spatiotemporal brain dynamics. We then demonstrated that this is due to ability of CHARM to capture the critical, non-local/long-range nature of brain dynamics, by established the underlying mechanisms through using a precise whole-brain model. Importantly, we also demonstrated that CHARM reveals the significantly different critical dynamics of wakefulness and sleep in the manifold network interactions. Taken together, CHARM is a promising theoretical framework for capturing the complex network dynamics in neuroscience.

Overall, the results suggest that the low-dimensional manifold networks revealed by CHARM could explain how the human brain is able to solve complex computational problems despite the relative slowness of neuronal communication. Moreover, it is important to note that we use the mathematical structure of Schrödinger’s equation (which is particularly suited for expressing non-locality) to allow for the discovery of the low-dimensional manifolds of empirical brain dynamics. In contrast to the single neuron doctrine arising from the debates between Nobel Prize winners Ramón y Cajal and Golgi (Lopez-Munoz *et al*., 2006), this strongly suggests that brain computation does not primarily happen locally, but that computation is mainly a long-range network effect. As such, the results demonstrate the key causal role of manifold networks as a fundamental organising principle of brain function at the whole-brain scale, providing evidence that networks of brain regions rather than individual brain regions are the key computational engines of critical brain dynamics.

## Methods

### Empirical data acquisition and preprocessing: Human Connectome Project

Resting state data was acquired from the Human Connectome Project described below in details.

#### Ethics

The Washington University–University of Minnesota (WU-Minn HCP) Consortium obtained full informed consent from all participants, and research procedures and ethical guidelines were followed in accordance with Washington University institutional review board approval (Mapping the Human Connectome: Structure, Function, and Heritability; IRB # 201204036).

#### Participants

The data set used for this investigation was selected from the March 2017 public data release from the Human Connectome Project (HCP) where we chose a sample of 1003 participants, all of whom have resting state data.

#### Neuroimaging acquisition for fMRI HCP

The 1003 HCP participants were scanned on a 3-T connectome-Skyra scanner (Siemens). We used one resting state fMRI acquisition of approximately 15 minutes acquired on the same day, with eyes open with relaxed fixation on a projected bright cross-hair on a dark background as well as data from the seven tasks. The HCP website (http://www.humanconnectome.org/) provides the full details of participants, the acquisition protocol and preprocessing of the data for both resting state and the seven tasks. Below we have briefly summarised these.

#### Neuroimaging acquisition for fMRI HCP

The 1003 HCP participants were scanned on a 3-T connectome-Skyra scanner (Siemens). We used one resting state fMRI acquisition of approximately 15 minutes acquired on the same day, with eyes open with relaxed fixation on a projected bright cross-hair on a dark background as well as data from the seven tasks. The HCP website (http://www.humanconnectome.org/) provides the full details of participants, the acquisition protocol and pre-processing of the data for both resting state and the seven tasks. Below we have briefly summarised these.

The pre-processing of the HCP resting state and task datasets is described in details on the HCP website. Briefly, the data is pre-processed using the HCP pipeline which is using standardized methods using FSL (FMRIB Software Library), FreeSurfer, and the Connectome Workbench software (Glasser *et al*., 2013; Smith *et al*., 2013). This standard pre-processing included correction for spatial and gradient distortions and head motion, intensity normalization and bias field removal, registration to the T1 weighted structural image, transformation to the 2mm Montreal Neurological Institute (MNI) space, and using the FIX artefact removal procedure (Navarro Schroder *et al*., 2015; Smith *et al*., 2013). The head motion parameters were regressed out and structured artefacts were removed by ICA+FIX processing (Independent Component Analysis followed by FMRIB’s ICA-based X-noiseifier (Griffanti *et al*., 2014; Salimi-Khorshidi *et al*., 2014)). Pre-processed timeseries of all grayordinates are in HCP CIFTI grayordinates standard space and available in the surface-based CIFTI file for each participants for resting state and each of the seven tasks.

We used a custom-made Matlab script using the ft_read_cifti function (Fieldtrip toolbox (Oostenveld *et al*., 2011)) to extract the average timeseries of all the grayordinates in each region of the Mindboggle-modified Desikan-Killiany parcellation (Desikan *et al*., 2006) with a total of 62 cortical regions (31 regions per hemisphere) (Klein and Tourville, 2012), which are defined in the HCP CIFTI grayordinates standard space. The BOLD timeseries were filtered using a second-order Butterworth filter in the range of 0.008-0.08Hz.

### Empirical data acquisition and pre-processing: Human sleep data

#### Ethics

Written informed consent was obtained, and the study was approved by the ethics committee of the Faculty of Medicine at the Goethe University of Frankfurt, Germany.

#### Participants

We used fMRI- and PSG data from 18 participants taken from a larger database that reached all four stages of PSG (Stevner *et al*., 2019; Tagliazucchi and Laufs, 2014). Exclusion criteria focussed on the quality of the concomitant acquisition of EEG, EMG, fMRI, and physiological recordings.

#### Acquisition and pre-processing of fMRI and polysomnography data

Neuroimaging fMRI was acquired on a 3 T system (Siemens Trio, Erlangen, Germany) with the following settings: 1505 volumes of T2*-weighted echo planar images with a repetition time (TR) of 2.08 seconds, and an echo time of 30 ms; matrix 64 x 64, voxel size 3 x 3 x 2 mm^3^, distance factor 50%, FOV 192 mm^2^.

The EPI data were realigned, normalised to MNI space, and spatially smoothed using a Gaussian kernel of 8 mm^3^ FWHM in SPM8 (http://www.fil.ion.ucl.ac.uk/spm/). Spatial downsampling was then performed to a 4 x 4 x 4 mm resolution. From the simultaneously recorded ECG and respiration, cardiac- and respiratory-induced noise components were estimated using the RETROICOR method (Glover *et al*., 2000), and together with motion parameters these were regressed out of the signals. The data were temporally band-pass filtered in the range 0.008-0.08 Hz using a sixth-order Butterworth filter. We extracted the timeseries in the DK62 parcellation (Tzourio-Mazoyer *et al*., 2002).

Simultaneous PSG was performed through the recording of EEG, EMG, ECG, EOG, pulse oximetry, and respiration. EEG was recorded using a cap (modified BrainCapMR, Easycap, Herrsching, Germany) with 30 channels, of which the FCz electrode was used as reference. The sampling rate of the EEG was 5 kHz, and a low-pass filter was applied at 250 Hz. MRI and pulse artefact correction were applied based on the average artefact subtraction method(Allen *et al*., 1998) in Vision Analyzer2 (Brain Products, Germany). EMG was collected with chin and tibial derivations, and as the ECG and EOG recorded bipolarly at a sampling rate of 5 kHz with a low-pass filter at 1 kHz. Pulse oximetry was collected using the Trio scanner, and respiration with MR-compatible devices (BrainAmp MR+, BrainAmp ExG; Brain Products, Gilching, Germany).

Participants were instructed to lie still in the scanner with their eyes closed and relax. Sleep classification was performed by a sleep expert based on the EEG recordings in accordance with the AASM criteria (2007). Results using the same data and the same pre-processing has previously been reported (Stevner *et al*., 2019; Tagliazucchi and Laufs, 2014).

### Neuroimaging structural connectivity and extraction of functional timeseries

#### Parcellations

All neuroimaging data was processed using the DK80 parcellation. This consists of the Mindboggle-modified Desikan-Killiany parcellation (Desikan *et al*., 2006) with a total of 62 cortical regions (31 regions per hemisphere) (Klein and Tourville, 2012), as well as well 18 subcortical regions (nine regions per hemisphere): hippocampus, amygdala, subthalamic nucleus (STN), globus pallidus internal segment (GPi), globus pallidus external segment (GPe), putamen, caudate, nucleus accumbens and thalamus. This created a parcellation with 80 regions in the DK80 parcellation; also precisely defined in the HCP CIFTI grayordinates standard space with a total of 91,282 grayordinates (sampled at 2 mm^3^).

### Empirical data: Diffusion MRI for tractography

#### Neuroimaging acquisition for dMRI HCP

We obtained multi-shell diffusion-weighted imaging data from 985 subjects of the HCP 1200 data release. The standard acquisition protocol takes 59 minutes (six runs of each approximately 9 minutes and 50 seconds). We also obtained diffusion spectrum and T2-weighted imaging data from 32 participants from the HCP database who were scanned for a full 89 minutes. The acquisition parameters for both groups are described in details on the HCP website (Setsompop *et al*., 2013).

#### Generating structural connectivity matrices from dMRI

In order to be as precise as possible for the model fitting, we estimated the structural connectivity matrix from two HCP dMRI datasets. The first dataset, Standard HCP dMRI, uses the highest quality multi-shell diffusion data acquired in sequence taking 59 minutes from 985 HCP participants (HCP data acquired at Washington University in St. Louis) (Li *et al*., 2019; Van Essen *et al*., 2013) (see HCP specifications on their website). The second dataset, Special HCP dMRI, uses even better protocols taking 89 minutes for each of 32 HCP participants at the MGH centre. Both dMRI datasets were preprocessed and made available as part of the freely available Lead-DBS software package (http://www.lead-dbs.org/).

The precise preprocessing is described in details in Horn and colleagues (Horn *et al*., 2017), but briefly, the data was processed using a generalized q-sampling imaging algorithm implemented in DSI studio (http://dsi-studio.labsolver.org). Segmentation of the T2-weighted anatomical images produced a white-matter mask and co-registering the images to the b0 image of the diffusion data using SPM12. In each HCP participant, 200,000 fibres were sampled within the white-matter mask. Fibres were transformed into MNI space using Lead-DBS (Horn and Blankenburg, 2016). The methods used the algorithms for false-positive fibres shown to be optimal in recent open challenges (Maier-Hein *et al*., 2017; Schilling *et al*., 2019). The risk of false positive tractography was reduced in several ways. Most importantly, this used the tracking method achieving the highest (92%) valid connection score among 96 methods submitted from 20 different research groups in a recent open competition (Maier-Hein *et al*., 2017). We subsequently used the standardized methods in Lead-DBS to produce the structural connectomes for the DK80 parcellation used in the whole-brain Hopf model.

### Theoretical methods

#### Reconstruction in the source fMRI Space for the generalization dataset

In order to test the ability of a given manifold to reconstruct the source signal on a generalisation set of data, we need to solve the “pre-image” problem. Let us define a diffusion map (Gaussian or Complex) on a training set. The generalisation set can be defined by 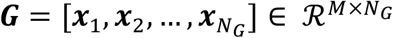, given by *N*_*G*_ time points of BOLD signal of the *M* brain regions that were not included in the training set used for the definition of the manifold space. We can then measure the ability of manifolds generated by different methods of reconstructing and generalising the source data.

#### Reconstruction using harmonic decomposition methods

Explicit inverse mappings from manifold to source space are not well-defined for non-linear manifold learning algorithms (such as harmonic decomposition and CHARM) (Evangelou *et al*., 2023; Papaioannou *et al*., 2022; Patsatzis *et al*., 2023). Most often, the lifting of predictions made on the manifold back to the source space problem is solved by using the Nyström extension methodology (Nyström, 1929), which has been derived from the solution of the Fredholm integral equation of the second kind. Here, we adopted this widely used framework (Evangelou *et al*., 2023; Papaioannou *et al*., 2022; Patsatzis *et al*., 2023), which can be described by:

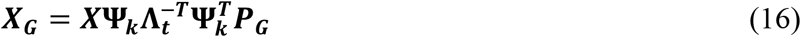

where **Ψ**_***k***_ = [***φ***_1_, ***φ***_2_, …, ***φ***_*k*_] is the matrix of the first *k* eigenvectors spanning the reduced manifold space. For the case of Gaussian diffusion, **Λ**_***t***_ is the corresponding diagonal matrix for the *k* first eigenvalues at *t*-time diffused steps, i.e. the diagonal elements are [λ_0_^*t*^, λ_1_^*t*^, …, λ_*k*_^*t*^].

For the case of CHARM, **Λ**_***t***_ is the diagonal matrix for the *k* first eigenvalues of the transition matrix 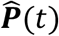, i.e. the diagonal elements are 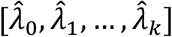. The matrix 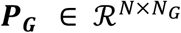 is given by the first *N* rows and the last *N*_*G*_ columns of the matrix 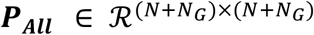 defined by the *t*-time diffused steps transition matrix for all data points included in the training and generalisation set. This matrix ***P***_***All***_ is computed with the corresponding for each case, affinity matrix for all data points, that is for the harmonic decomposition: 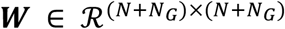 (for the each single time step diffusion) and for CHARM: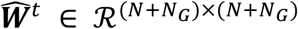. In other words, as before ***P***_***All***_ belongs to the weighted graph of the data set, but now including all training and generalisation data points, that is *N* + *N*_*G*_ time points observation of the BOLD signal of the *M* brain regions.

#### Quantitative measurements characterising the manifold reduction

For comparing the quality of the manifold reduction obtained by the two frameworks (harmonic decomposition and CHARM), we compute the 1) reconstruction error, 2) edge metastability and 3) conservation of edge metastability between source and manifold spaces.

#### Reconstruction Error

We assess the quality of the manifold reduction in the latent space by comparing the empirical and reconstructed functional connectivity (FC) for each participant. Given the matrix BOLD signal observation ***X***, we define for each brain region *m*, the time series of observations ***s***_*m*_ by the *m*-row of ***X***. The empirical FC (***FC***^*emp*^) is computed by the Pearson correlation, that is the elements are given by

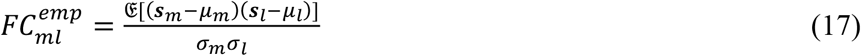

where for brain regions *m* and *l, μ*_*m*_ and *μ*_*l*_ are the corresponding mean values across time, and σ_?_ and σ_B_ are the corresponding standard deviations across time. In the equation, 𝔈[] denotes the expected value operator. Similarly, the FC for the reconstructed time series (***FC***^*rec*^) is computed by

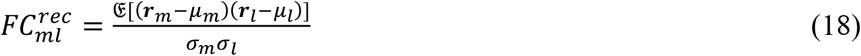

Where ***r***_*m*_ are the time series of reconstructed timeseries for the *m*-row of ***X***. We use two different metrics for comparing the functional connectivity. Let us denote by ***f***^*emp*^ and ***f***^*rec*^ the elements of the corresponding FC matrices. We compute the mean squared error for *M* regions

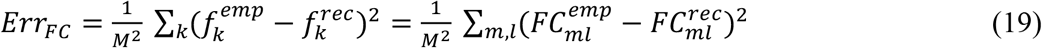

and the correlation between the elements of the FC matrices is given by

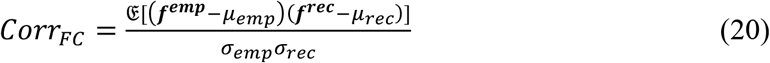

where *μ*_*emp*_,*μ*_*rec*_ are the corresponding mean values, and σ_*emp*,_σ_*rec*_ the corresponding standard deviations of the elements.

#### Edge-metastability in manifold space

The measure of edge-metastability was recently introduced as sensitive way of comparing matrices inspired by the spatiotemporal variability of edge time series, introduced to capture fine-scale dynamics in fMRI recordings (Faskowitz *et al*., 2020; Sporns *et al*., 2021; Zamani Esfahlani *et al*., 2020). The resulting edge time series are formed by a simple procedure involving 1) z-scoring each of the two nodal time series independently and 2) and forming an edge time series by taking the element-wise product of the z-scored time series. Values of the edge time series reflect the co-fluctuation pattern between nodes. A positive co-fluctuation results when, at a specific point in time, both series are concordant relative to each of their mean signals. A negative co-fluctuation value results when, at a specific point in time, one time series is above the mean (a positive value) and the other is below the mean (a negative value). Notably, the mean of an edge time series equals the Pearson correlation. Edge time series have the same temporal resolution as the original data, allowing for the analysis of instantaneous (i.e., a single time frame) co-fluctuation patterns. This data has the dimensionality of edge-by-time.

We used this framework to compute for each participant the edge-metastability, which as just shown is a measure of the variability of the co-fluctuation patterns. This is a generalisation of the functional connectivity dynamics but computed for a slicing window of only one time point. For a given participant, we define a spatiotemporal series 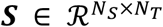 where the *N*_*S*_ rows correspond to space dimensions and the *N*_*T*_ columns to the timepoints. We note that in source space this matrix corresponds to ***X***, while in the manifold space corresponds to ***Y***. First, we z-score across the columns the time series for each space dimension, where the z-scored matrix ***S*** across its columns is called 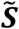 . The corresponding edge-centred matrix 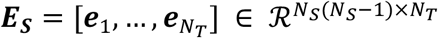 is defined as follows. Each column corresponds to a timepoint *t*, where the column is defined as a vector combining all pairwise combinations of the spatial dimension at time *t*. With *N*_*S*_ space dimensions, this results in *N*_*S*_(*N*_*S*_ − 1) pairs. From this matrix of pairs, we define the corresponding functional connectivity matrix 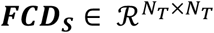,

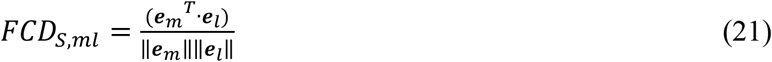

where the dot product denotes scalar product.

Let us denote by ***g***_*S*_ the upper diagonal elements of the ***FCD***_***S***_ matrix. Then, the edge-metastability of the spatiotemporal signal ***S*** (with variance 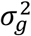) is given by the Gaussian entropy

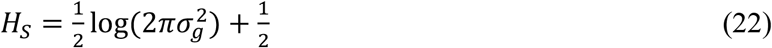

In order to measure the degree of spatiotemporal variability in latent space, we will use **Equation 22** for the spatiotemporal matrix ***Y*** in the different manifold spaces corresponding to the different framewoeks. We compute the edge-metastability in manifold space, *H*_*Y*_, for each participant.

#### Conservation of edge-metastability between source and manifold space

In order to quantify how well the empirical edge-metastability in source space is expressed in the different manifold spaces, we calculate the edge-metastability for each participant in source space, i.e. *H*_*X*_ and in manifold space *H*_*Y*_, and perform the correlation between these two measurements across participants. We use C_*XY*_ to denote this mass of conservation of metastability between source and manifold space.

#### Time asymmetry between networks in manifold space

We quantified the time asymmetry between the networks in the manifold by computing the shifted functional connectivity as follows:

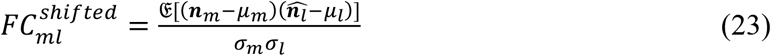

Where the time series of observations ***n***_*m*_ is the time series of the network *m*, and 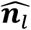 is the time series of network *i* shifted one TR (2.08 s) forward in time. The corresponding mean values across time are *μ*_*m*_ and *μ*_*l*_; σ_*m*_ and σ_*l*_ stand for the corresponding standard deviations across time. In the equation, 𝔈[] denotes the expected value operator.

### Whole-Brain Model

In order to demonstrate the effects of long-range functional correlations and of rare anatomical long-range connections, we used whole-brain modelling which links anatomical structural connectivity with functional dynamics through a model of local dynamics in each brain region (Breakspear, 2017; Deco and Kringelbach, 2014; Deco *et al*., 2015; Kringelbach and Deco, 2020). The anatomical structural connectivity (SC) is determined *in vivo* using diffusion MRI (dMRI) in conjunction with probabilistic tractography. The whole-brain model creates a suitable balance between complexity and realism by using the connectivity between brain regions (reflected in SC) to reproduce the empirically-measured whole-brain dynamics included those measured with fMRI (Breakspear, 2017). Such whole-brain models have had widespread success in explaining the brain dynamics of many different brain states (Breakspear, 2004; Deco et al., 2011; Deco et al., 2013; Deco and Kringelbach, 2014; Ghosh et al., 2008; Honey et al., 2007).

Here, the local dynamics of each brain region are modelled using a Stuart-Landau oscillator, which is equivalent to the normal form of a supercritical Hopf bifurcation. This is a powerful model for examining the shift from noisy to oscillatory dynamics (Kuznetsov, 1998). Whole-brain Hopf models have been able to replicate key aspects of brain dynamics observed in electrophysiology (Freyer *et al*., 2011; Freyer *et al*., 2012), magnetoencephalography (Deco *et al*., 2017a) and fMRI (Deco *et al*., 2019; Kringelbach *et al*., 2020). Specifically, the whole-brain dynamics of neuroimaging data with timeseries from a parcellation of total *M* regions can be expressed by coupling the local dynamics of *M* Stuart-Landau oscillators coupled via the connectivity matrix ***C***

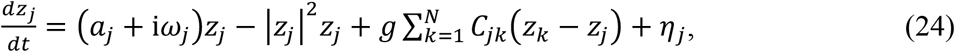

where for the oscillator in region *j*, the complex variable *z*_*j*_ denotes the state (*z*_*j*_ = *x*_*j*_ + *iy*_*j*_), *η*_*j*_ is additive uncorrelated Gaussian noise with variance σ^2^ (for all *j*), *ω*_*j*_ is the intrinsic node frequency, and *a*_*j*_ is the node’s bifurcation parameter. Within this model, the intrinsic frequency *ω*_*j*_ of each node is in the 0.008–0.08Hz band, which has been shown to be the optimal capturing neural dynamics. The intrinsic frequencies were estimated from the data, as given by the averaged peak frequency of the narrowband blood-oxygen-level-dependent (BOLD) signals of each brain region. The global coupling parameter is denoted by *g*. For *a*_*j*_ *>* 0, the local dynamics settle into a stable limit cycle, producing self-sustained oscillations with frequency *ω*_*j*_/(2π). For *a*_*j*_ < 0, the local dynamics present a stable spiral point, producing damped or noisy oscillations in the absence or presence of noise, respectively. The fMRI signals were modelled by the real part of the state variables, i.e., *x*_*j*_ = Real(*z*_*j*_).

It has been shown that the best working point for fitting whole-brain neuroimaging dynamics is at the brink of the bifurcation, that is with *a*_*j*_ slightly negative but very near to zero (usually *a*_*j*_ = −0.02) (Deco *et al*., 2017b). In the results, we show that for fitting the HCP resting data, the position where the coupling parameter achieve the best fitting is also critical, in the sense that is maximising the variability across time of the Kuramoto order parameter (also called Kuramoto metastability). The Kuramoto order parameter is a measure of synchronization between the different brain regions, and is given by:

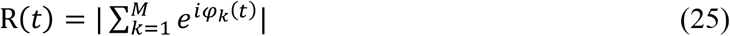

where *ϕ*_*k*_(*t*) are the phases of the spatiotemporal data (empirical or simulations), extracted by the classical Hilbert transform. The Kuramoto metastability is defined by the variance of R across time. The maximal value of R obtained at the optimal fitting point can be used as a proxy of criticality, given that the variability of R can be thought of as the susceptibility. It is well-known that a maximum of susceptibility corresponds to a trace criticality and consequently provides a meaningful measure of the relevance of long-range functional correlations.

### Support vector machine for classification

We used machine learning for both pattern separation and classification by using a support vector machine (SVM) with Gaussian kernels as implemented in the Matlab function *fitcecoc*. The function returns a full, trained, two class, error-correcting output codes (ECOC) model. This is achieved using the predictors in the input with class labels. The function uses K(K – 1)/2 binary SVM models using the one-versus-one coding design, where we used K=2 as the number of unique class labels. The output was two classes corresponding to the conditions (after versus before, or responder versus non-responder, or psilocybin versus escitalopram treatment). The input features used for classification were shifted functional connectivity (for each participant and condition). We trained the SVM with the leave-one-out cross-validation procedure, i.e. by randomly choosing one patient for generalisation and the whole rest for training, repeated and shuffled 1,000 times. Furthermore, the training set was balanced in terms of number of examples for each class label, and randomly selecting the participants in each class for each shuffling iteration.

## Notes

### Competing Interest Statement

The authors have declared no competing interest.

